# Polyploidy promotes transformation of epithelial cells into non-professional phagocytes

**DOI:** 10.1101/2025.03.24.645044

**Authors:** Yi-Chun Huang, Caique Almeida Machado Costa, Nicolas Vergara Ruiz, Xianfeng Wang, Allison Jevitt, Christina Marie Breneman, Chun Han, Wu-Min Deng

## Abstract

Removal of dead and damaged cells is critical for organismal health. Under stress conditions such as nutritional deprivation, infection, or temperature shift, the clearance of nonessential cells becomes a universal strategy to conserve energy and maintain tissue homeostasis. Typically, this task is performed by professional phagocytes such as macrophages. However, non-professional phagocytes (NPPs) can also adopt a phagocytic fate under specific circumstances. Similar to professional phagocytes, NPPs undergo transitions from immature to mature states and activation, but the precise cellular and molecular mechanisms governing their maturation, induction and phagocytic execution remain largely unknown. A notable example of stress-induced phagocytosis is the removal of germline cells by follicle cell-derived NPPs during oogenesis in *Drosophila*. In this study, we report that the transformation of follicle cells into NPPs is dependent on Notch signaling activation during mid-oogenesis. Moreover, Notch overactivation is sufficient to trigger germline cell death and clearance (GDAC). We further show that polyploidy, driven by Notch signaling-induced endoreplication, is essential for the transformation of follicle cells into NPPs. Polyploidy facilitates the activation of JNK signaling, which is crucial for the phagocytic behavior of these cells. Additionally, we show that polyploidy in epidermal cells, another type of NPPs, is important for their engulfment of dendrites during induced degeneration. Together, these findings suggest that polyploidy is a critical factor in the transformation of epithelial cells into NPPs, enabling their phagocytic functions, which are essential for maintaining cellular and organismal homeostasis during stress conditions.

**SIGNIFICANCE:** The ability to remove dead and damaged cells is essential for maintaining tissue homeostasis and organismal health. While this task is typically performed by professional phagocytes such as macrophages, non-professional phagocytes (NPPs) can also acquire phagocytic functions during development or in response to stress conditions. Using *Drosophila* oogenesis as a model, we reveal that the transformation of follicle cells into NPPs is driven by Notch signaling and is critically dependent on polyploidy. Our findings show that polyploidy, induced through Notch signaling-mediated endoreplication, is required for activating JNK signaling, a pathway essential for the phagocytic behavior of NPPs. Furthermore, we show that polyploidy also facilitates the phagocytic activity of epidermal cells in clearing degenerating dendrites. Together, these results suggest that polyploidy plays an important role in enabling epithelial cells to adopt NPP functions and in maintaining tissue and organismal homeostasis under stress conditions.

## Introduction

Tissue homeostasis relies on a delicate balance between cell survival and death, and the prompt clearance of dead cells is critical, as failure to remove them can lead to chronic inflammation and various disorders. The clearance of dead cells is typically performed by professional phagocytes, such as macrophages and neutrophils, through a process called efferocytosis. Additionally, non-professional phagocytes (NPPs), which generally do not engage in phagocytosis under normal conditions, can also adapt to perform phagocytic functions when confronted with cellular debris, particularly in contexts where professional phagocytes are unavailable. This adaptive capability ensures a flexible response to cell death, promoting the prompt removal of cellular debris to maintain tissue homeostasis and prevent the development of pathological conditions (1-7).

NPPs are widely used during development to remove germline cells, neurons and secretory cells. Germline-cell death and clearance (GDAC) is a conserved developmental process in metazoans (8-13). For example, mammalian spermatogenesis relies heavily on the role of NPPs, such as Sertoli cells in testes (10, 14). Located within the seminiferous tubules of the testes, Sertoli cells support sperm development and remove excess germ cells by engulfing and clearing apoptotic cells and cellular debris through phagocytosis, thereby contributing to the maturation of male germ cells(10, 11). Granulosa cells, found in ovarian follicles, play a pivotal role in folliculogenesis by supporting the development and maturation of ovarian follicles(15, 16). Recently, programmed cell death and extracellular acidification have been observed to facilitate the removal of nurse cells by pregranulosa cells during oocyte development in prenatal mouse embryos(17, 18), indicating a potentially conserved mechanism for ensuring germ cell quality across species(19). Mature granulosa cells have also been identified as NPPs that clear dying granulosa cells and oocytes during follicle atresia(16).

*Drosophila* oogenesis relies on developmental checkpoints to respond to environmental and physiological stresses, ensuring the quality of the mature egg(3, 20). Germline cells are enveloped by somatically derived epithelial cells known as follicle cells (FCs), which provide essential nutrients, signals, and protection to nurture oocyte growth(21-24). Recent research has unveiled another pivotal function of follicle cells as NPPs. Under biotic and abiotic stresses such as starvation, cold acclimation, infection or drug administration (e.g., rapamycin), impaired oogenesis leads to the death of germline cells during the middle stages of oogenesis (stage 8-9) in female flies(25-28). In response, follicle cells transform into NPPs, engaging in the phagocytosis of deceased germline cell debris. The mechanisms underlying this transition, from the normal developmental functions of the follicle cells to their role in stress-induced emergency management, remains incompletely understood. This transformation requires activation of the Jun N-terminal Kinase (JNK) signaling pathway, the phagocytic receptor Draper (Drpr), and the modulator of cytoskeletal changes and GTPase Rac1, as well as CED-12/ELMO, which facilitate cytoplasmic expansion and the engulfment of dead nurse cell debris(18, 29, 30).

In *Drosophila*, other NPPs such as glial cells and epidermal cells contribute significantly to maintaining tissue integrity of the nervous system during development and stress. Glial cells play a crucial role in the nervous system by engulfing apoptotic neurons and axonal debris, a process dependent on the Drpr and the JNK signaling pathway(31, 32). This process ensures proper neural pruning and repair mechanisms. Similarly, epithelial cells in the larval epidermis clear pruned and injured sensory dendrites, facilitating tissue remodeling and immune responses. Both glial and epithelial NPPs rely on JNK signaling to regulate the cytoskeletal dynamics necessary for engulfment and on phagocytic receptors like Draper and Ced-12 to recognize and internalize dying cells(18, 31, 32).

Here we report that NPP transformation in follicle cells depends on a Notch-induced differentiation and maturation process. Previously, we have shown that the differentiation of follicle cells, accompanied with a mitotic to endocycle (ME) switch during mid-oogenesis (stages 7-9), is dependent on activation of Notch signaling in follicle cells(33, 34). At the onset of ME switch, Delta (Dl), a Notch ligand, is upregulated in the germline cells to induce canonical Notch signaling in the follicle cell epithelium(34, 35). In this study, we show that Notch-induced polyploidy is essential for the maturation of NPPs. Polyploidy allows activation of the JNK signaling in follicle cells, resulting in morphological changes of the cells for effective phagocytosis. Similarly, larval epidermal NPPs requires polyploidy for efficient clearance of degenerating sensory dendrites, corroborating the important roles of endoduplication in the maturation of epithelial NPPs.

## Results

### 1. The temporal regulation of stress-induced germline cell death and clearance (GDAC)

To investigate the temporal pattern of stress-induced germline cell death and clearance (GDAC), we exposed adult female flies to various stressors, including starvation, γ-ray irradiation, *Pseudomonas entomophila* (*Pe*) infection, and tumor implants in the visceral cavity(18, 36-38) (Fig. 1A-E). The ovaries of these stressed female flies consistently exhibited the GDAC phenotype during mid-oogenesis (Fig. 1B-F, Supplemental Fig. 1A-A””), indicating a general temporal regulation of GDAC during oogenesis.

**Figure 1:**
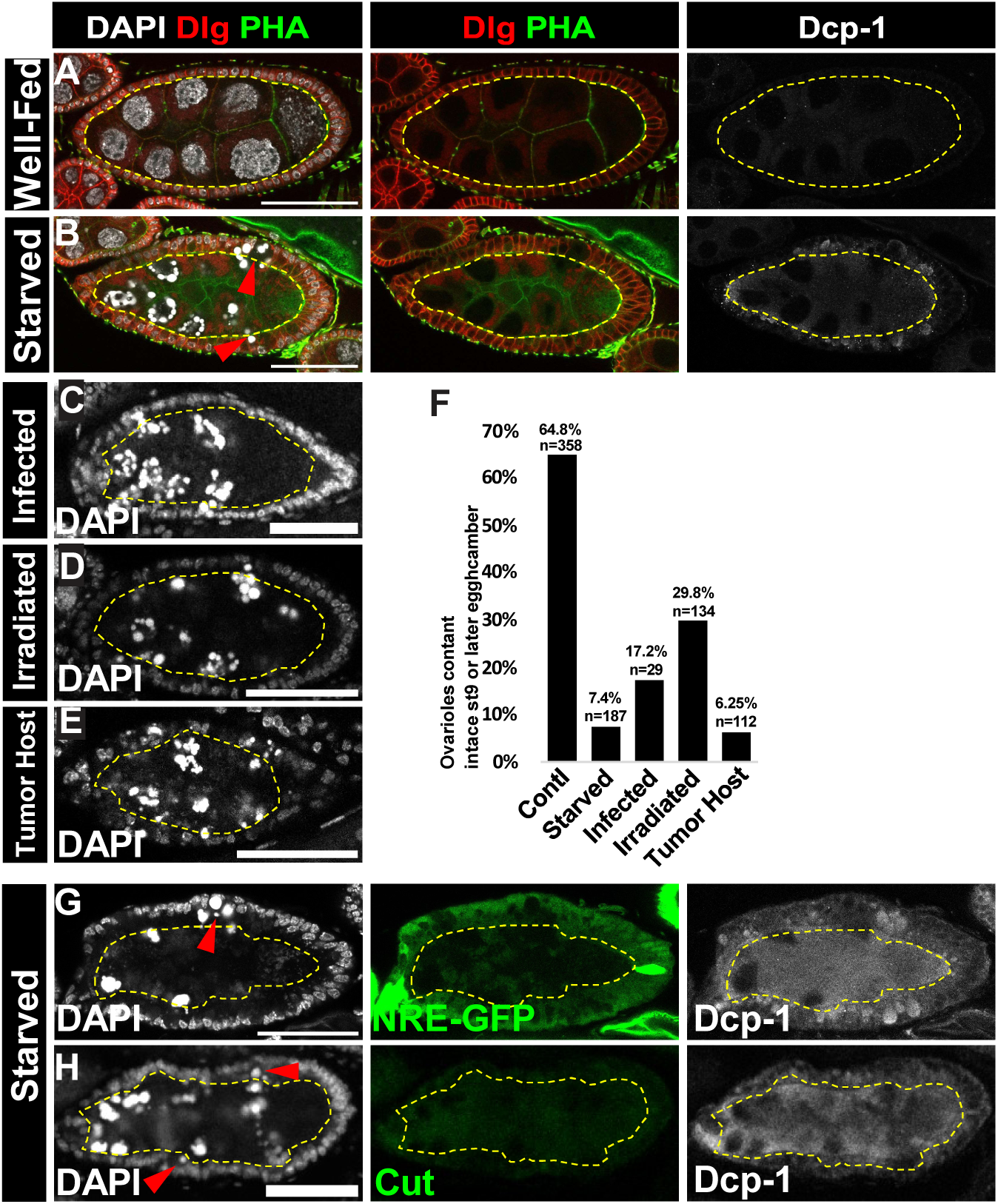
Stress-induced germline cell death and clearance (GDAC). (A,B) Confocal images comparing (A) well-fed and (B) starved stage (st) 8 egg chambers. Dlg labels the lateral membrane of follicle cells (FCs) and phalloidin (PHA) detects F-actin. Dead germline debris caused by starvation were engulfed by follicle cells (FCs). Dcp-1 serves as the Apoptosis marker. (C-E) GDAC egg chambers were observed in (C) *Pe*-infected, (D) Gamma-ray-irradiated, and (E) tumor-implanted flies. (F) Quantitative analysis of intact stage-9 or later egg chambers per ovarioles under different stress conditions. (G, H) Confocal images showing that GDAC occurred in middle-stage egg chambers, indicated by higher NRE-GFP levels and the absence of Cut expression. Red arrowheads indicate dying germline debris engulfed by FCs. Dashed yellow circles indicate the germline area. DAPI marks cell nuclei. Posterior is oriented to the right. Scale bars: 50 μm.

Since follicle cell differentiation and maturation depends on Notch signaling(34, 39), we explored whether there is a correlation between Notch activation and stress-induced GDAC. To this end, we examined the expression of the Notch-responsive element (NRE-GFP), a reporter of Notch activity(40), in egg chambers undergoing starvation-induced GDAC. Our findings revealed that GDAC occurred in egg chambers where Notch signaling was activated in the FCs (Fig. 1G). Furthermore, we stained these egg chambers with an antibody against Cut, a homeodomain protein and marker of immature follicle cell fate that is negatively regulated by Notch signaling(39). We observed an absence of Cut expression in egg chambers undergoing GDAC (Fig. 1H). These results suggest that GDAC occurs in egg chambers when Notch signaling is activated in follicle cells during mid-oogenesis (Fig. 1G-H, Supplemental Fig. 1B).

### 2. Notch signaling is required in follicle cells for stress-induced GDAC

To determine whether Notch signaling activation is necessary for stress-induced GDAC, we disrupted Notch signaling in follicle cells by knocking down *mastermind* (*mam*), a transcriptional co-activator of the Notch pathway(41), using a pan-follicle cell Gal4 driver (*traffic jam (tj)-Gal4)*(42). The *tj>mam^IR^* intervention impaired GDAC induced by starvation, leaving uncleared remnants of germline cell nuclei in degenerating egg chambers (Fig. 2A, B, Supplemental Fig. 1C). To quantify the GDAC phenotype, we measured the ratio of the germline area to the total egg chamber area as an indicator of germline clearance efficiency (Supplemental Fig. 1D) (18). Compared to control starved egg chambers, the germline area was significantly larger when *mam* expression was knocked down in follicle cells (Fig. 2C). These results indicate that phagocytosis by follicle cells was compromised due to reduced Notch signaling, underscoring the essential role of Notch signaling in follicle cell-mediated GDAC.

**Figure 2:**
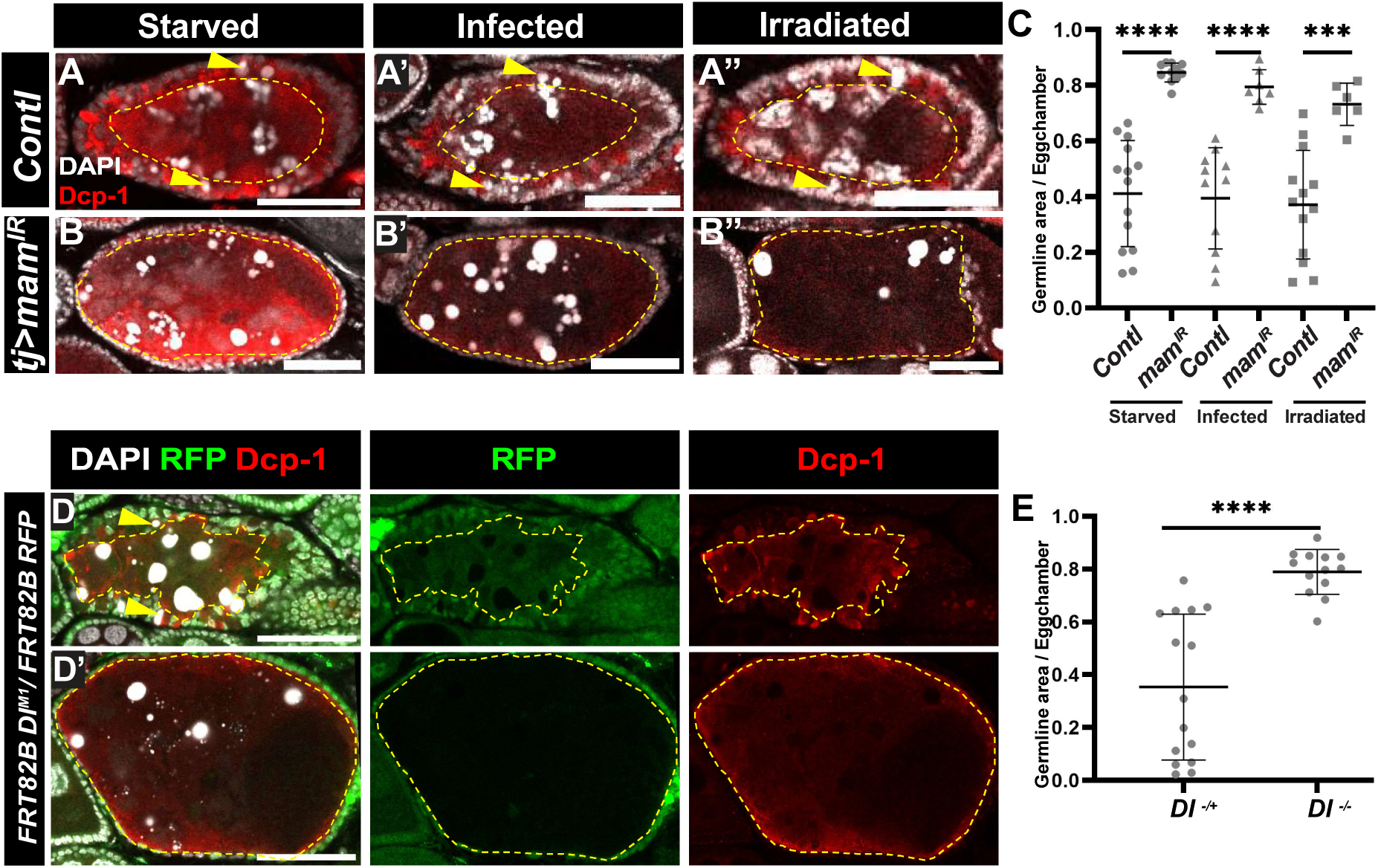
Notch signaling is required for stress-induced GDAC. (A,B) Confocal images comparing stress-induced GDAC in (A-A”) wild-type and (B-B”) *mam* knock-downed egg chambers. (C) Graph comparing germline areas between wild-type and Notch-signaling deficient egg chambers. The Y-axis represents the percentage of germline area relative to follicle cell area, and the X-axis indicates genotypes. (D, D’) Confocal images show (D) a starved wild-type egg chamber with normal GDAC and (D’) a mosaic egg chamber with a *Dl*-mutant germline clone (no GFP expressed in the germline area, indicated by the yellow dashed cycle), resulting in reduced engulfment. (E) Graph comparing germline areas between egg chambers of wild-type and *Dl* mutant germline clones. The Y-axis represents the percentage of germline area relative to entire egg chamber area, and the X-axis indicates genotypes. **** represents p-value ≤ 0.0001, *** indicates p-value ≤ 0.001. Posterior is oriented to the right. Scale bars: 50 μm.

To explore the involvement of Notch signaling in follicle-cell NPPs under other stress conditions, we inoculated flies with the *Pe* or exposed them to γ -radiation (Fig. 2A’, A”). Consistently, when *mam* was knocked down in follicle cells, the clearance of dying germline cells was hindered (Fig. 2B’, B” & C). These findings suggest that Notch and its downstream transcriptional activation are critical for GDAC induced by stressors, further demonstrating the requirement of Notch activation in follicle cells for effective NPP function.

Notch activation in follicle cells during mid-oogenesis is driven by the upregulation of the Delta (Dl) ligand in germline cells(34). To determine whether germline-expressed Dl is necessary for starvation-induced GDAC, we generated mosaic egg chambers containing *Dl* germline clones and observed that starvation-induced engulfment of dying germline cells was blocked (Fig. 2D’). Despite this, apoptosis in germline cells still occurred, as indicated by fragmented nuclei and elevated expression of Dcp-1, an apoptosis marker(43). In contrast, in egg chambers heterozygous for *Dl* in the germline, engulfment of dying germline cells was preserved, leading to a reduced germline area (Fig. 2D, E, & Supplemental Fig 2E). These findings suggest that Notch signaling, activated by germline-derived *Dl*, is crucial for the transformation of follicle cells into NPPs during stress-induced GDAC.

### 3. Notch hyperactivation as a novel stress to induce GDAC

The finding that Notch signaling is necessary for follicle cells to transform into NPPs prompted us to investigate the effects of Notch hyperactivation in these cells. Using the *tj-Gal4^ts^* driver to overexpress the Notch intracellular domain (NICD), the active form of Notch, in follicle cells, we observed that oogenesis was arrested during stages 8-9 (Fig. 3A). The arrested egg chambers exhibited dying germline cells with reduced germline area (Fig. 3A, dashed yellow line), as revealed by elevated expression of the apoptosis marker Dcp-1 (Fig. 3B) and positive TUNEL staining(43) (Fig. 3C). Examination of these degenerating egg chambers revealed that the dying germline cells were engulfed by the follicle cells, similar to stress-induced GDAC (Fig. 1B-E). Germline cytoplasm, marked by the germline marker Vasa, and condensed nuclear DNA of nurse cells were enclosed by elongated and expanded follicle cells (Fig. 3E). The germline area continued to diminish and was eventually cleared, leaving follicle cells behind (Supplemental Fig. 2A). These observations indicate that the follicle cells, upon Notch hyperactivation, had transformed into NPPs and phagocytosed the dead germline cell debris.

**Figure 3:**
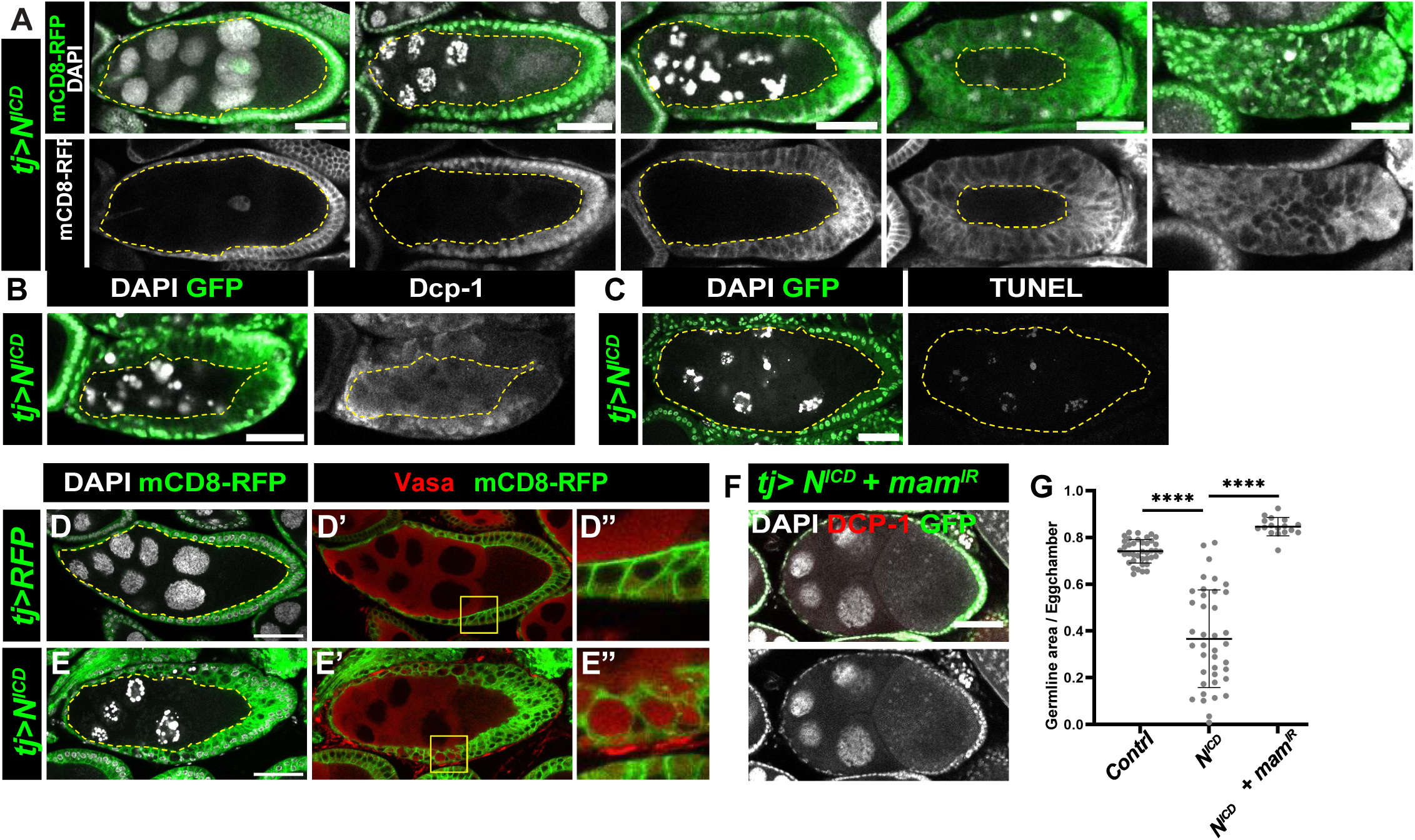
Hyperactivation of Notch signaling in follicles cell induces GDAC. (A) Progression of NICD-overexpression induced GDAC in stages 8 and 9 egg chambers. Germline nuclei degenerated and were engulfed by follicle cells (FCs), resulting in the germline area (yellow dashed cycle) shrinking and eventually disappearing. (B, C) Cell death markers Dcp-1 (B) and TUNEL (C) appeared in egg chambers undergoing NICD-induced GDAC. (D-D”) In control egg chambers, Vasa was restricted to the germline area. (E-E”) in NICD-induced GDAC egg chamber, Vasa was detected in areas encircled by FCs, indicating engulfment (white arrows). (F) Co-knockdown of *mam* (the co-activator of the Notch pathway) blocked NICD-induced GDAC. The round shape of the egg chamber displayed a typical phenotype associated with defective Notch signaling. (G) Graph comparing germline areas in st 8/9 egg chambers between wild-type, NICD overexpression, and NICD overexpression with *mam* knockdown. The Y-axis represents the percentage of germline area relative to follicle cells area, The X-axis indicates genotypes. **** represents p-value ≤ 0.0001, *** represents p-value ≤ 0.001. Dashed yellow circles indicate the germline area. DAPI (white) marks cell nuclei. Posterior is oriented to the right. Scale bars: 50 μm.

To determine whether NICD-overexpression (OE)-induced GDAC is dependent on transcriptional activation, we simultaneously knocked down *mam*, the co-activator, in NICD-OE follicle cells. This intervention sufficiently suppressed the GDAC phenotype (Fig 3F, G), indicating that Notch-mediated transcriptional regulation is essential for the transformation of follicle cells into NPPs.

Together, these findings suggest that Notch hyperactivation functions as a stress-like signal to induce GDAC, resembling the conditions observed in egg chambers under starvation, infection, and other stresses.

### 4. Notch hyperactivation-induced GDAC depends on phagocytic genes and JNK activation

In *Drosophila*, engulfment of dying germline cells depends on the phagocytic machinery genes *Draper* (*Drpr*), *Ced-12,* and *Rac1*(18, 44). To determine whether Notch hyperactivation-induced NPP transformation also relies on these genes, we knocked down *Drpr*, *Ced-12*, or *Rac1*, in follicle cells with NICD overexpression. Although the nurse cell nuclei were fragmented, indicating cell death, the germline area failed to diminish, suggesting the clearance of dying germline cells was impaired in all three cases (Fig 4 A-D, Supplemental Fig 2C). These findings indicate that NICD-OE-induced GDAC uses the established phagocytic pathway.

**Figure 4:**
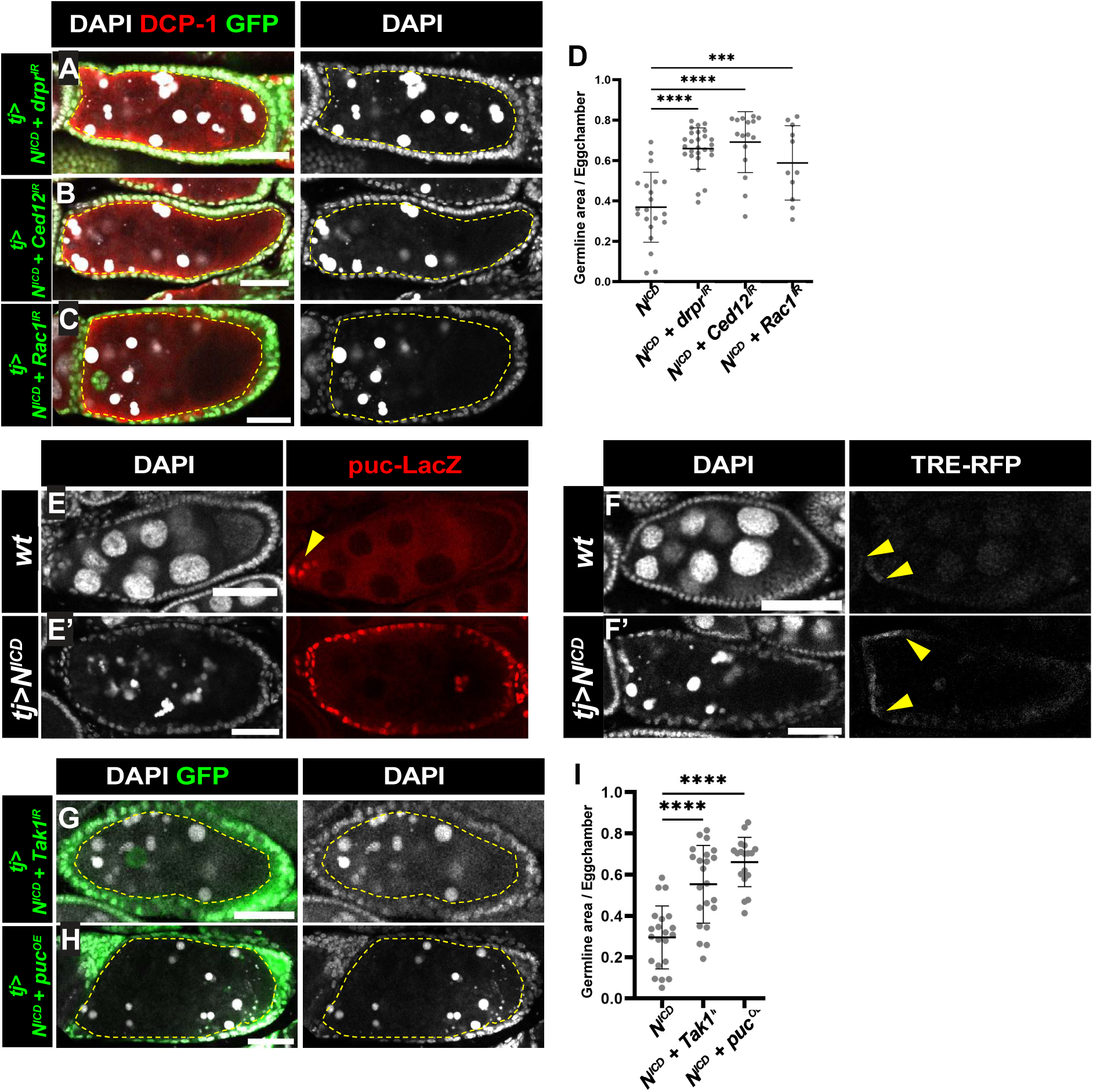
Notch-hyperactivation induced GDAC depends on phagocytic genes and JNK activation. (A-C) Confocal images showing the reduced engulfment ability of NPPs when phagocytic genes were knocked down in Notch-hyperactivation FCs. Dcp-1 (red) marks apoptosis. (A) Knockdown of *drpr*, (B) Knockdown of *Ced12,* and (C) Knockdown of *Rac1*. (D) Graph comparing germline areas with GDAC between NICD-overexpression (OE) along and NICD-OE with knockdown of *drpr*, *Ced12*, or *Rac1*, respectively. (E) The JNK reporter *puc-Lacz* was expressed only in the anterior follicle cells (yellow arrowhead) of a wild-type st 8/9 egg chamber. (E’) In an egg chamber with NICD-induced GDAC, *puc-Lacz* was expressed in all FCs. (F) In a wild-type st 8/9 egg chamber, another JNK reporter, TRE-RFP, was observed in anterior FCs. (F’) In egg chambers with NICD-induced GDAC was detected in a larger number of FCs. (G, H) Confocal images showing the GDAC was reduced by (G) knockdown of *Tak1* (a JNKKK) and (H) overexpression of *puc* (a negative regulator of JNK). (I) Graph comparing germline areas in egg chamber with GDAC between NICD-OE along and NICD-OE with *Tak1* knockdown or *puc*-OE, respectively. (A-C & G-H) Dashed yellow cycles indicate the germline area. (D & I) The Y-axis represents the percentage of germline area relative to the follicle cell area, and the X-axis indicates genotypes. **** represents p-value ≤ 0.0001, *** represents p-value ≤ 0.001. DAPI (white) marks cell nuclei. Posterior is oriented to the right. Scale bars: 50 μm.

During starvation-induced GDAC, JNK signaling is activated in follicle cells that are transformed into the NPPs(18). This activation is critical for engulfment behavior, and Drpr has been identified as a JNK target(8). To determine whether NPP transformation induced by Notch hyperactivation also depends on JNK signaling activation, we examined the expression of *puckered* (*puc*)*-LacZ*, a JNK reporter(45), in NICD-OE egg chambers. *puc* encodes a serine/threonine protein phosphatase that negatively regulates the JNK pathway(46). NICD overexpression led to cell-autonomous upregulation of *puc-LacZ* in follicle cells in egg chambers undergoing GDAC (Fig. 4E). Additionally, a TRE-RFP reporter, another JNK signaling reporter(47), was upregulated in Notch hyperactivation-induced NPPs (Fig 4F & Supplemental Fig 2D).

Next, we tested whether JNK signaling activation is required for the phagocytic behavior induced by NICD overexpression, similar to starvation-induced GDAC(48). To this end, we inhibited JNK signaling in NICD-OE follicle cells by knocking down *TGF-β activated kinase 1 (Tak1),* a positive regulator of the JNK pathway(49), and by misexpressing *puc*, a negative regulator of JNK. In both cases, the phagocytosis phenotype was alleviated (Fig. 4G-I, Supplemental Fig 2C).

Together, these findings demonstrate that JNK signaling activation is critical for Notch-hyperactivation induced phagocytic functions.

### 5. Polyploidization is a crucial event in NPP transformation in follicle cells

The pivotal role of Notch signaling in inducing the ME switch in follicle cells to become polyploid(50, 51) led us to hypothesize that polyploidy is a prerequisite for their transformation into NPPs. To test this, we employed the *tj-Gal4^ts^* driver to knock down *fizzy-related* (*fzr*), a known inducer of the endocycle(52), thereby blocking the ME switch. These flies were subsequently starved, and the *tj>fzr^IR^* egg chambers exhibited compromised GDAC (Fig 5A-C, Supplemental Fig 3A). Notably, some egg chambers retained nurse cells with normal nuclear morphology but exhibited fewer FCs (Supplemental Fig 3B), suggesting their viability is compromised. We also observed that reducing follicle cell polyploidy disrupted developmental germline removal (Supplemental 3C), which takes place during stages 13-14 of oogenesis removing the remnants of nurse cells by stretched follicle cells(53, 54). Similarly, in NICD-induced GDAC, knocking down *fzr* also resulted in decreased efficiency in germline corpse clearance (Fig 5D-F, supplemental 3D).

**Figure 5:**
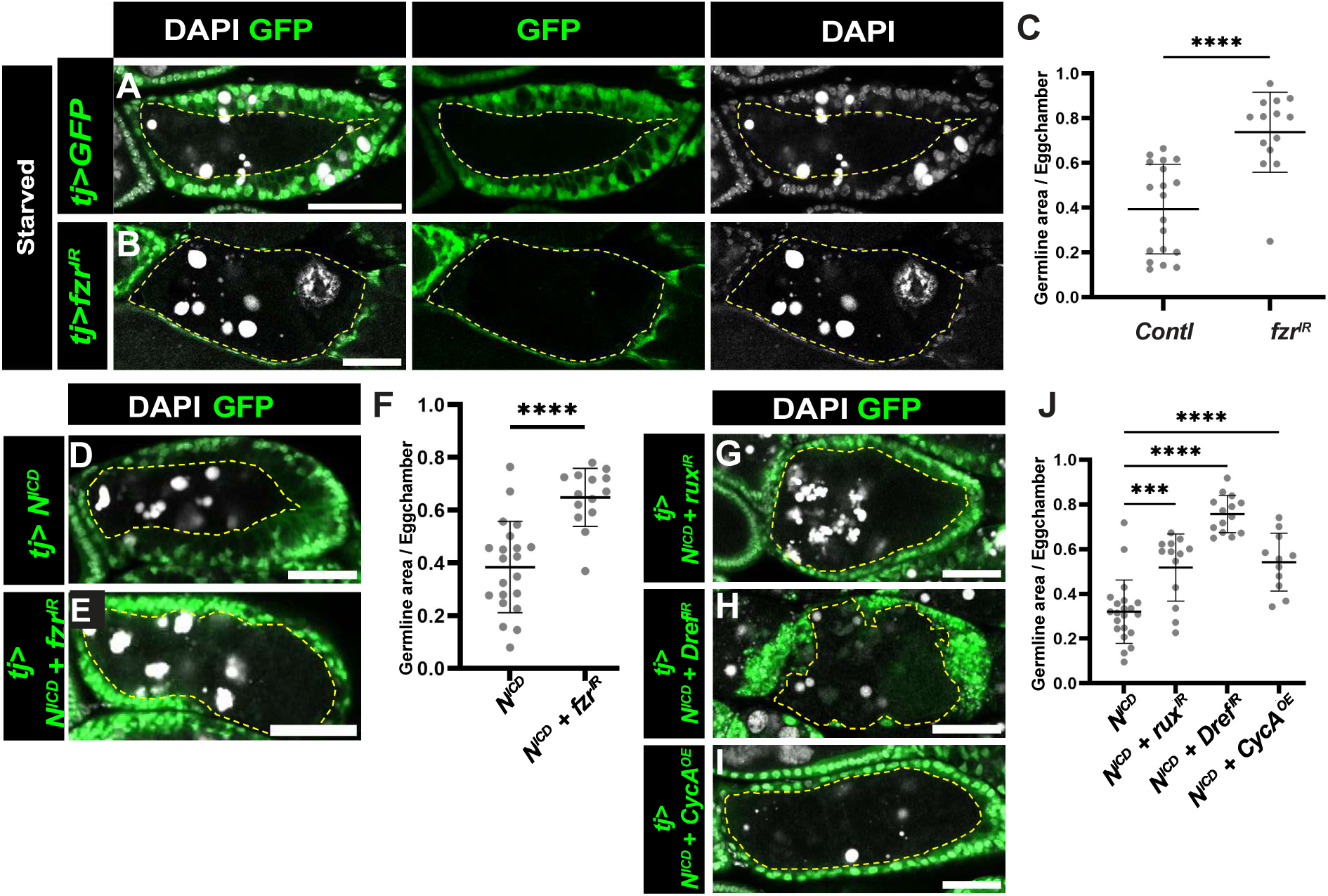
Polyploidy is a key downstream event for Notch-induced maturation of NPPs. (A,B) Confocal images show that during starvation, FCs in the control egg chambers (A) can engulf dying germline cells; however, (B) reducing FCs ploidy by knockdown of *fzr* impaired their effectiveness in cleaning dying germline debris. (C) Graph comparing germline areas in egg chambers with GDAC between control (*tj>GFP*) and *tj>fzr^IR^*. (D, E) Confocal images show that reducing FC ploidy by knocking down *fzr* in the NICD-OE background reduces the ability of FCs to clean the germline area debris. (F) Graph comparing the germline areas in egg chambers with GDAC between NICD-OE alone and NICD-OE with *fzr* knockdown. (G—I) Confocal images show that in NICD-OE background, reducing FC ploidy by (G) knocking down *rux* (a negative regulator of CycA), (H) knocking down *Dref* (The DNA replication-related element-binding factor), or (I) overexpressing CycA blocks effective transformation of FCs into NPPs. (J) Graph comparing the germline areas in egg chambers with GDAC between NICD-OE alone and NICD-OE with knockdown of *rux,* or *Dref,* or CycA-OE, respectively. (A, B, D, E, G-I) Dashed yellow cycles indicate germline area. (C, F, & J) The Y-axis represents the percentage of germline area relative to follicle cells area, and the X-axis indicates genotypes. **** represents p-value ≤ 0.0001, *** represents p-value ≤ 0.001. DAPI (white) marks cell nuclei. Posterior is oriented to the right. Scale bars: 50 μm.

To further validate these findings, we used an alternative strategy to inhibit endoreplication by targeting *roughex* (*rux*), a regulator of the cell cycle that inhibits Cyclin A (CycA)(55). Consistent with the results from *fzr* knockdown, interference with *rux* further demonstrated the critical role of polyploidy in enabling effective GDAC (Fig 5 G, J). Moreover, knocking down *Dref* or overexpression of *CycA*, which also disrupted endoreplication, blocked infection-induced GDAC (Fig 5 H-J), reinforcing the link between polyploidy and phagocytic capability. In addition, we tested the effect of *Dref* knockdown in infection-induced GDAC and observed impaired engulfment of dying germline cells (Supplemental Fig 3E).

These findings collectively suggest that endoreplication and subsequent polyploidization in follicle cells are critical for their transformation into NPPs. The development of polyploidy appears to equip these cells with the cellular machinery to meet the demands associated with GDAC under stress conditions.

### 6. Polyploid cells exhibit high levels of JNK signaling activity during stress-induced NPP transformation

To understand the differences between polyploid and diploid follicle cells in their ability to become NPPs, we generated mosaic egg chamber with *Notch* knockdown using the heat-shock inducible flip-out Gal4 to drive *Notch* RNAi expression. Such egg chambers are sensitive to stresses as degenerative egg chambers were frequently observed. As the follicular epithelium (FE) in these mosaic egg chambers contained both polyploid and diploid follicle cells in the same egg chamber, we could monitor their different response and behavior when facing germline cell death.

In these mosaic egg chambers, polyploid wild-type cells, which are positive for the Hindsight (Hnt, aka. Peb) signal—a downstream target of Notch signaling in follicle cells (33)— exhibited high levels of TRE-GFP (or TRE-RFP)(47), the JNK reporter (Fig 6A & Supplemental Fig 4A). By contrast, the *Notch*-knockdown diploid cells showed no or low levels of JNK activity (Fig 6A & Supplemental Fig 4A). Similarly, during stress-induced GDAC, knockdown of *Dref* in follicle cells had a similar effect on the expression of the JNK activity reporter. Polyploid wildtype follicle cells exhibited higher upregulation of the JNK reporter, whereas *Dref*-knockdown cells, marked by RFP, showed low levels of JNK activity (Supplemental Fig 4B, C). These findings suggest that polyploid cells are more prone to stress-induced JNK activation, which is crucial for NPP transformation.

**Figure 6:**
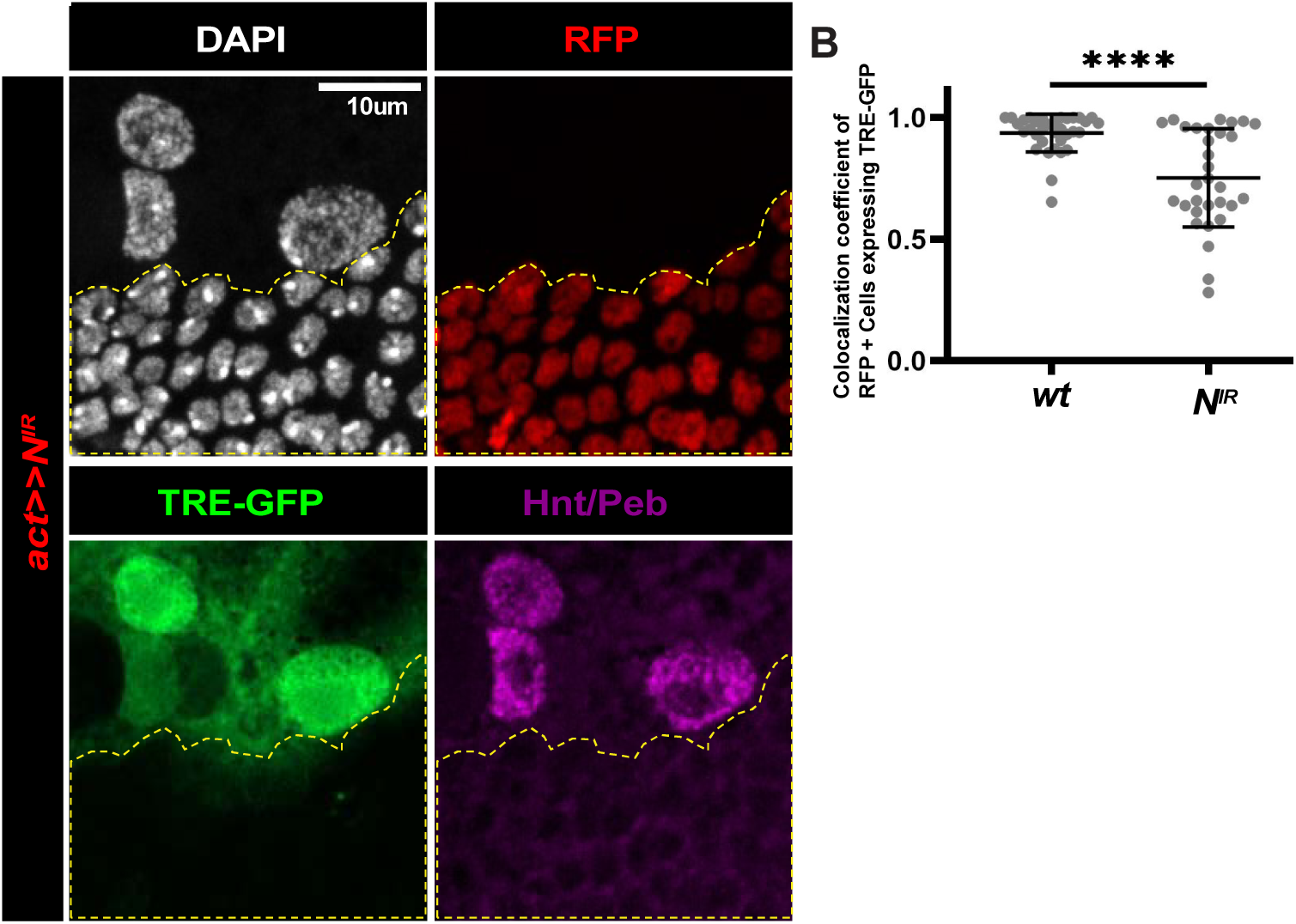
Polyploid follicle cells show high levels of JNK activation during NPP transformation. (A) Confocal images of mosaic follicular epithelium (FE) with Flip-out *Notch*-knockdown clones. Polyploid wild-type FCs (Hnt-positive, magenta) exhibited strong TRE-GFP expression, indicating high levels of JNK activity. In contrast, *Notch*-knockdown diploid FCs (RFP-positive, dashed yellow cycle) lacked TRE-GFP expression, indicating an absence of JNK activation in these cells. (B) Graph showing the colocalization coefficient between the RFP+ cells (cells with *Notch*-knockdown) and TRE-GFP, the JNK activity reporter. The coefficient ranges from 0 to 1, where 0 indicates no colocalization and 1 reflects complete colocalization across all pixels in both channels. The Y-axis represents the colocalization coefficient of RFP+ cells expressing TRE-GFP, and the X-axis indicates genotypes. **** represents p-value ≤ 0.0001, DAPI (white) marks cell nuclei. Scale bars: 10 μm.

### 7. Polyploidy of epidermal cells is important for their phagocytosis of degenerating dendrites

To further explore the relationship between polyploidy and NPP transformation, we examined the effects of inhibiting endoreplication on larval epidermal cells. These polyploid epidermal cells are NPPs and can efficiently engulf degenerating sensory dendrites that are normally attached to them(56). To inhibit epidermal endoreplication, we knocked down *Dref* using two distinct RNAi lines via *R16D01-Gal4*, which is expressed in a stripe of epidermal cells in the middle of each segment(56). These manipulations resulted in 62% reduction in cell size for both RNAi lines (Fig. 7A-D) in 3^rd^ instar larvae, confirming the importance of endoreplication in epidermal cell growth. To examine whether these reductions in cell size correlates with reduced phagocytic capabilities, we then tested knockdown of *Dref* in the context of genetically induced neurodegeneration via the knockout of nicotinamide mononucleotide adenyltransferase (Nmnat) in class IV dendritic arborization (C4da) neurons(57). C4da neurons grow elaborate dendritic arbors, which can be visualized by the dendritic marker *ppk-MApHS* (Fig. 7E). Nmnat is an enzyme required for the synthesis nicotinamide adenine dinucleotide (NAD+), whose knockout in C4da neurons results in phagocytosis-dependent dendrite degeneration in late third instar larvae(57), as reflected by greatly reduced dendrites and dendrite debris dispersed throughout epidermal cells (Fig. 7F, F’). Knocking down *Dref* from the posterior part of each segment via *en-Gal4* resulted in much longer dendrites and much less dendrite debris in the *en-Gal4* region in comparison to the *Nmnat* knockout alone (Fig.7F-J), indicating impaired engulfment by epidermal cells. Altogether, these data suggest that endoreplication, resulting in polyploidy, is important for the phagocytic capabilities of epidermal cells.

**Figure 7:**
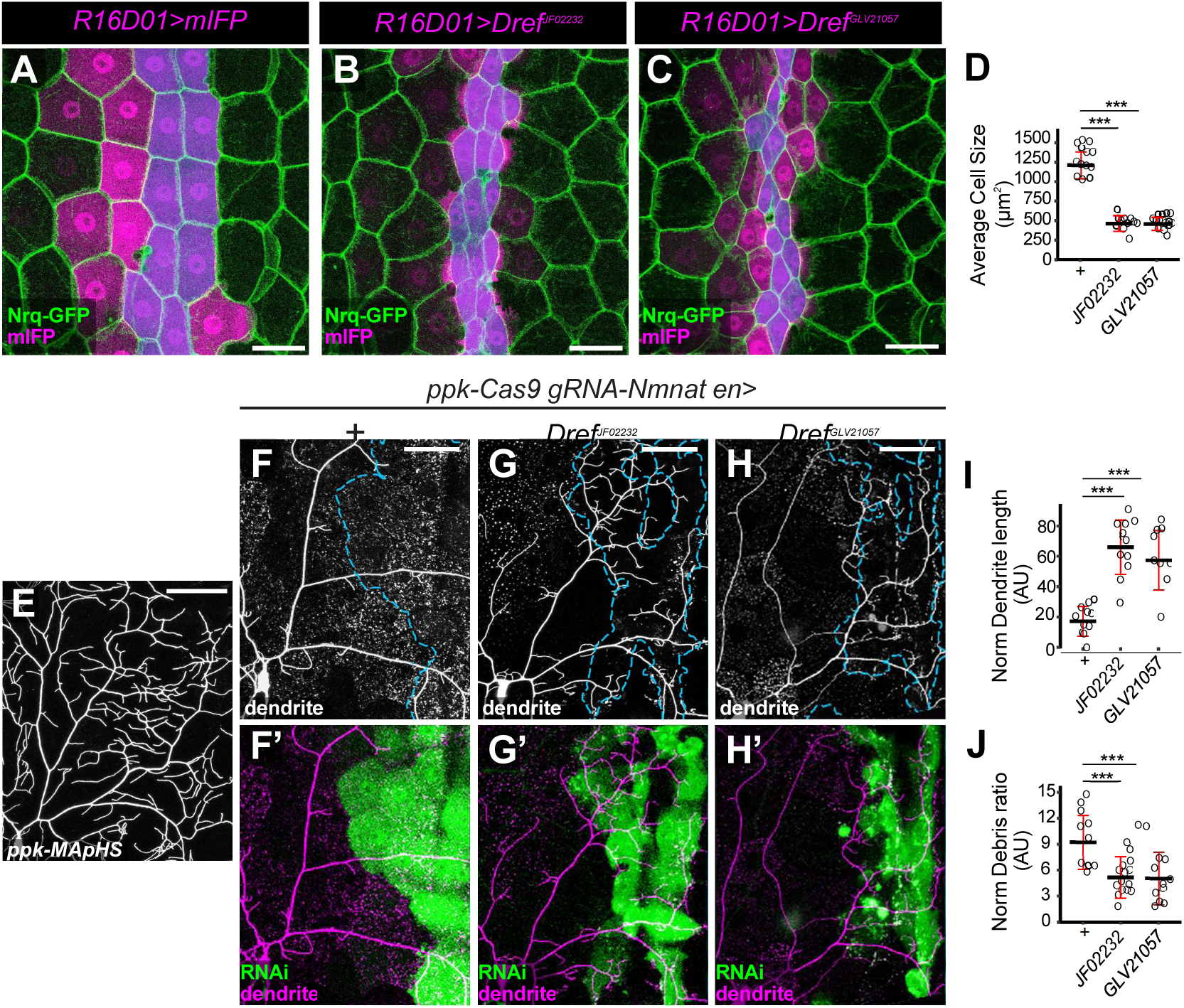
Polyploidy of epidermal cells is important for their phagocytosis of degenerating dendrites. (A-C) Epidermal cells in *R16D01-Gal4* control (A) and *R16D01-Gal4*-driven *Dref* RNAi (JF02232 in B and GLV21057 in C) expressing animals. The *R16D01-Gal4* domain is labeled by UAS-mIFP expression (magenta). Epidermal cell borders are labeled by Nrg-GFP (green). The blue overlay indicates the measured Gal4-expressing epidermal cells. (D) Average epidermal cell size in the *R16D01-Gal4* region. n = number of neurons and N = number of animals: control (n = 20, N = 5); JF02232 (n = 22, N = 7); GLV21057 (n = 25, N = 5). (E) A WT class IV dendritic arborization (C4da) neuron labelled by the *tdTom* channel of *ppk-MApHS*. (F-H’) Dendrites of *Nmnat*-deficient C4da neurons (via *ppk-Cas9*-driven knockout of *Nmnat*) in *en-Gal4* control (F-F’), JF02232 (G-G’), and GLV21057 (H-H’) expressing animals. The merged images (F’, G’ H’) show neurons (*tdTom* channel of *ppk-MApHS*) in magenta and the *Gal4*^en^ domain (*UAS-mBFP*) in green. The *en-Gal4* domains in dendrite images are enclosed by cyan dashed lines. (I-J) Normalized dendrite length (I) and normalized debris ratio (J) in F-H. n = number of neurons and N = number of animals: control (n = 12, N = 5); JF02232 (n = 19, N = 8); GLV21057 (n = 14, N = 6). Scale bars: 50μm for all images. For all quantifications, ***p<0.001; one-way ANOVA with Tukey post-hoc test.

## Discussion

In this study, we show that polyploidy is a prerequisite for NPP transformation in two different contexts, including the clearance of dying germline cells in *Drosophila* egg chambers and the removal of degenerating dendrites by epidermal cells. In ovarian follicle cells, disrupting endoreplication by knocking down *fzr*, *rux*, or *Dref* impaired stress-induced GDAC, highlighting the critical role of polyploidy in these processes. Similarly, in polyploid epidermal cells that engulf degenerating dendrites, knockdown of *Dref* led to reduced cell size and impaired dendrite engulfment. These findings indicate a conserved requirement for polyploidy in NPP-mediated phagocytosis across different tissues. Supporting this broader principle, glial cells, another class of NPPs, are also known to undergo polyploidization during development, reinforcing the importance of polyploidy in NPP functionality(58, 59).

Follicle cells, with their clear temporal-spatial pattern of development, offer an excellent model for understanding the crucial events and signaling pathways that prepare them to become NPPs(3, 5, 27, 34). The maturation of follicle cells is induced by Notch signaling, which plays an essential role in coordinating the development between germline and somatic cells(23, 60). Our previous studies showed that Notch activation in follicle cells trigger a mitotic to endocycle (ME) switch at stages 6/7 of oogenesis(34). In this study, we confirmed that early follicle cells are incapable of performing phagocytic functions(8, 61), and showed that the temporal regulation of follicle cell maturation by Notch signaling is critical for NPP maturation. The canonical Notch signaling pathway, along with downstream components like the transcriptional co-activator Mam, is critical for enabling follicle cells to efficiently phagocytose dead cells. These findings suggest that the maturation of NPPs through Notch signaling is a prerequisite to NPP functionality, drawing parallels to the development of professional phagocytes. Professional phagocytes like macrophages differentiate from monocyte precursors, undergoing extensive proliferation and maturation before acquiring phagocytic capabilities. Their precursors in the mammalian system, monocytes, cannot phagocytose before differentiation(62) Notch signaling plays a vital role in this differentiation process, underscoring its conserved importance in both *Drosophila* and mammals(63, 64). This transition from precursor to fully functional phagocyte observed in *Drosophila* follicle cells closely mirrors the development of macrophages(65). The regulatory mechanisms governing follicle-cell maturation in *Drosophila* may provide valuable insights into how signaling pathways drive the differentiation and functional specialization of phagocytes in higher organisms.

JNK signaling plays a crucial role downstream of Notch signaling in follicle cell NPPs. Both starvation- and Notch hyperactivation-induced GDAC required JNK activation, as demonstrated by the upregulation of *puc*-LacZ and TRE reporters in follicle cells. Inhibition of JNK signaling through *Tak1* knockdown, or *puc* overexpression, significantly impaired GDAC, underscoring its necessity for phagocytic functions. Our study shows that mature, differentiated follicle cells activate the JNK signaling pathway, a process critical for their phagocytic transformation. This activation is most likely related to these cells being polyploid, an important event induced by Notch signaling in follicle cells(34, 35). While JNK activation was detectable in diploid follicle cells, it was significantly weaker. In contrast, polyploid cells exhibit stronger JNK activation, enabling them to acquire the cytoskeletal and molecular machinery necessary for phagocytosis. The differential responses to JNK signaling between polyploid and diploid cells may be the key distinction underlying their potential in NPP transformation.

Polyploidy has also been observed in professional phagocytes. In mammals, polyploidization through cell fusion is essential for the phagocytic function of osteoclasts, a specialized phagocyte involved in bone resorption(66). This process enhances osteoclast functionality by increasing their cellular machinery and metabolic capacity, enabling efficient bone degradation. Although the mechanisms of polyploidization in osteoclasts differ from those in *Drosophila* follicle or epidermal cells, the outcomes are strikingly similar. In both cases, polyploidization leads to the cessation of the mitotic cycle and specialized adaptations for phagocytosis. These parallels suggest that, despite differing mechanisms, polyploidy serves a conserved role in enabling cells to meet the demands of their specialized functions, such as phagocytosis.

In the context of oogenesis, our findings raise an important question: why do follicle cells undergo polyploidization? While it has been suggested that polyploidy equips these cells to support the metabolic and nutritional demands of the growing oocyte(67), our results indicate that polyploidy may also serve as a preparatory mechanism, enabling follicle cells to respond to environmental stressors such as nutritional shortages, sudden temperature changes, or infections. Under stress conditions, polyploid follicle cells are primed to sacrifice germline cells, engulf and digest germline material, and provide essential nutrients to sustain the organism. This adaptive strategy is reminiscent of plants, where polyploidization at the organismal level often enhances stress tolerance and survival(68). These findings suggest that polyploidy may represent a conserved ancient mechanism across the tree of life, allowing organisms to withstand environmental challenges and maintain resilience.

## Materials and Methods

### Fly strains and genetics

*Drosophila* stocks used in this study are listed in the STAR★Methods section (Supplemental Table 1). *Drosophila* maintains and crosses detail are in supplemental documents.

### Environmental Stress induction experiments

Starvation experiment: Adult flies were cultured on yeast food for one day, then transferred to 1% agar for 1-2 days and harvesting ovaries.

Gamma (γ)-ray Irradiation: Adult flies were exposed to a single dose of γ-ray (18-36Gy) using a Gammacell 40 Cs-137 irradiator (MDS Nordion, Mississauga, ON) with a dose rate of 36 Gy/hr. Flies were dissected 24 hrs post-irradiation.

*Pseudomonas entomophila* (*Pe*) Infection: *Pe* was cultured overnight in Luria broth (LB) containing rifampicin (100 µg/mL) at 30°C. The bacterial solution was then diluted to an OD_600_ of 0.05 with 1X PBS. Flies were stabbed in the thorax using insect pins and dissected the next day(69).

Tumor transplantation: The transplantation procedure was performed as previously described(70, 71). Briefily, the *N^ICD^*-overexpression induced imaginal ring tumors from laeval salivary glands(72) were transplanted into *w^1118^* adult flies. The transplanted flies were cultured at 29°C for two-three days before harvesting ovaries.

### Immunohistochemistry and confocal imaging

Overies were dissected as previously described(73). More detail (including TUNEL assay) in supplementary.

### Epidermal cell quantification and imaging

Epidermal cells were quantified as described previously(56). Briefly, images of epidermal cells labeled by Nrg-GFP taken with a 20X objective were processed in ImageJ sequentially by Gaussian Blur (Sigma: 1), Auto Local Threshold (Phansalkar method, radius: 30), Particles4 (to remove isolated particles, Skeletonize (2D/3D) to mark epidermal cell borders. Regions of interest (ROIs) were manually drawn to encompass the epidermal cells for quantification by Analyze Particles.

For imaging epidermal cells and dendrites of class IV da neurons, third instar larvae at 100 hr AEL were anesthetized with isoflurane for two minutes and mounted in glycerol. The A2, A3, and A4 segments were imaged with a Leica SP8 confocal using a 40x oil objective. Quantification and statistical analysis details in supplementary.

## Acknowledgments

We thank the Bloomington *Drosophila* Stock Center, Vienna *Drosophila* Resource Center, Kyoto *Drosophila* Stock Center, Developmental Studies Hybridoma Bank, and fly research community (K. McCall, D. Bilder, and D. Bohmann) for providing fly lines and antibodies. We also extend our graditude to William Palmer, whose previous work on Notch signaling in follicle cells inspired this study, and other member of the Deng lab for discussions and contributions.

## Supplementary

### Supplementary Materials and Methods

#### Fly strains and genetics

*Drosophila* lines were maintained and crossed on standard BDSC cornmeal food (recipe available at https://bdsc.indiana.edu/information/recipes/bloomfood.html). Crosses containing the temperature-sensitive (ts) *Gal80* transgene were raised at 18°C. The adult progeny from these crosses were shifted to 29°C for three days to drive transgene expression before harvesting ovaries. For crosses to generate flip-out clones, flies were cultured at 25°C. The adult progeny were heat-shocked at 37°C for 5-10 mins, then transferred to 29°C for two days before dissection or shifted to a stress assay.

#### Immunohistochemistry and confocal imaging

The following antibodies were used: anti-Cut (DSHB, mouse, 1:15), anti-Dlg (DSHB, mouse, 1:50), anti-Drpr (DSHB, mouse, 1:50), anti-Vasa (DSHB, rat, 1:300), anti-Hnt/Peb (DSHB, mouse, 1:15), anti-Dcp-1(Cell Signaling, rabbit, 1:200), anti-β-Galactosidase (Promega, mouse, 1:500). Primary antibodies were incubated overnight at 4°C. Alexa Fluor conjugated secondary antibodies (ThermoFisher Scientific, details listed in STAR★Methods) (Supplemental Table 1) were used at 1:400 and incubated at room temperature for 2 hours. Nuclei were labeled with DAPI (ThermoFisher Scientific, 1:1000). Phalloidin (ThermoFisher Scientific, 1:50). Samples were mounted and imaged using Zeiss LSM 800 or LSM 980 Confocal Microscopes.

#### TUNEL assay

TUNEL (Terminal deoxynucleotidyl transferase dUTP nick end labeling) assay was performed using In the Situ Cell Death Detection Kit, TMR red (Roche) following the manufacturer’s protocol. Briefly, ovaries were fixed in 4% EM-grade formaldehyde (Electron Microscopy Sciences), washed with PBT (0.2% Triton X-100 in PBS), and incubated in 100mM sodium citrate at 65°C for 30min. After another wash with PBT, the tissues were stained using the TUNEL kit, followed by standard DAPI and mounting.

#### Quantification and statistical analysis

Data analysis was conducted using GraphPad Prism, excpet for the colocalization coefficient in Fig 6B. An unpaired t-test was used for two-sample comparisons, while one-way ANOVA with Dunnett’s multiple comparison test or One-way ANOVA with Tukey’s multiple comparison test was applied for multiple-sample comparisons. The colocalization coefficient for was calculated using the Zen (Zeiss, Gemary) colocalization function, and statistical differences were analyzed using PAST (PAleontological STatistics). A non-parametric ANOVA (Kruskal-Wallis) test was performed, followed by Dunn’s post-hoc test for multiple comparisons. Specific statistical approaches for each figure are indicated in the corresponding figure legend.

#### Quantification of dendrites and debris

The method to quantifying debris resulting from dendrite degeneration has been previously described(1). Briefly, the pHluorin channel in MApHS was used to generate a dendrite mask, and tdTomato channel was used to create a mask for both dendrites and debris. The pHluorin area was subtracted from dendrites + debris area (tdTomato) to isolate a mask containing only the debris signal. A region of interest (ROI) was manually drawn to include a quadrant of a ddaC neuron’s territory. The methods for tracing and measuring C4da neuron dendrite length have been previously described(2). The ROI area, total dendrite length and debris area were measured. Dendrite length and debris area were normalized by the ROI area.

## Supplementary Figure Legends

**Supplementary Figure 1:**
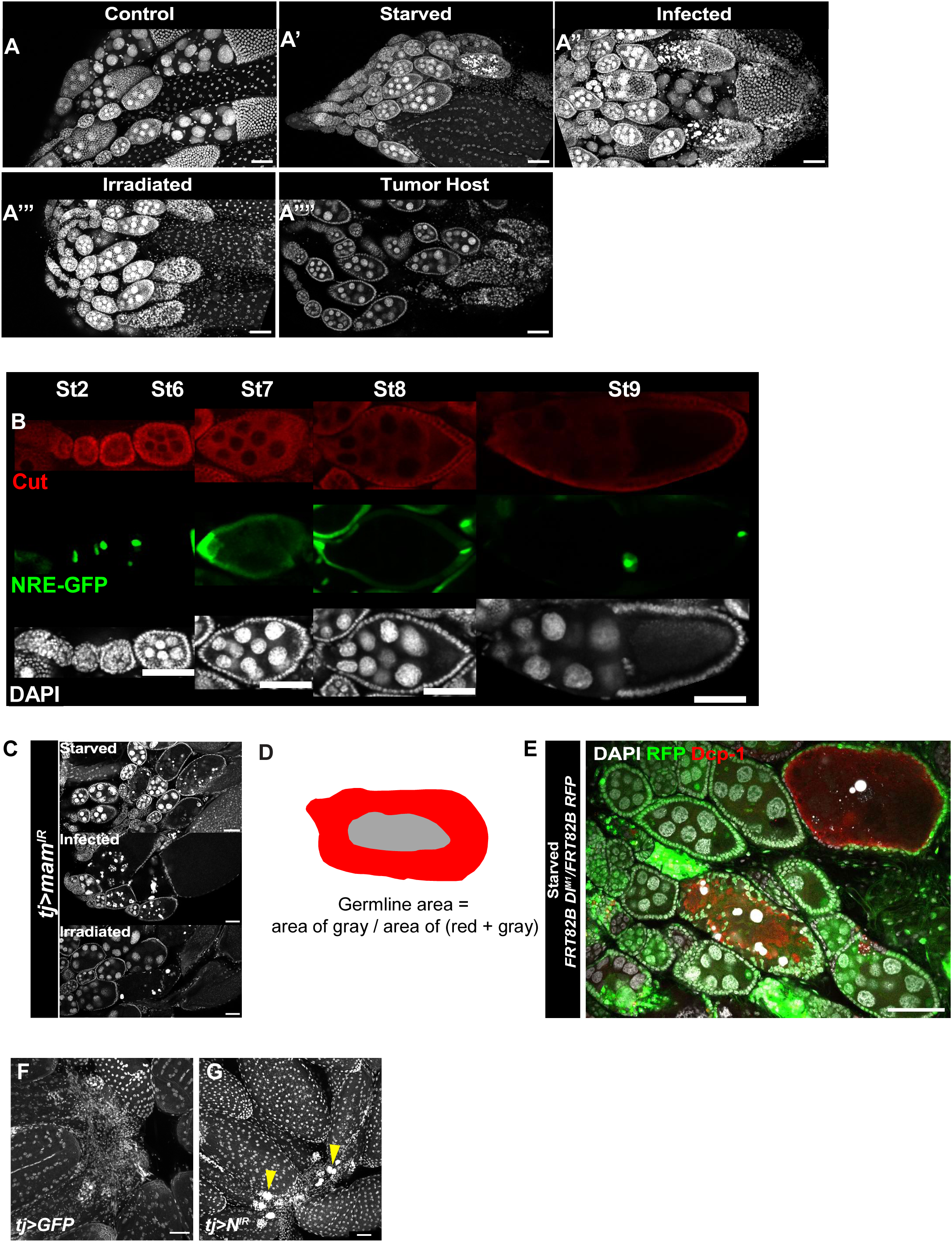
(A) Confocal images showing Cut expression restricted to main-body FCs during early oogenesis stage (st1-st6), while NRE-GFP, a Notch pathway reporter, begins expression after the mitotic-to-endocycle (ME) switch. (B) Wide-field confocal images corresponding to Figures1C-G. (C) Schematic illustration of how the germline area was calculated. (D) Wide-field confocal images of Figure 2B. (E) Wide-field confocal images of Figure 2D. (F-G) Confocal images of the corpus luteum / entrance to oviduct region. (F) Control fly. (G) N-knockdown flies show uncleaned germline debris (yellow arrowhead). DAPI was used to mark cell nuclei. Posterior is oriented to the right. Scale bars: 50 μm.

**Supplementary Figure 2:**
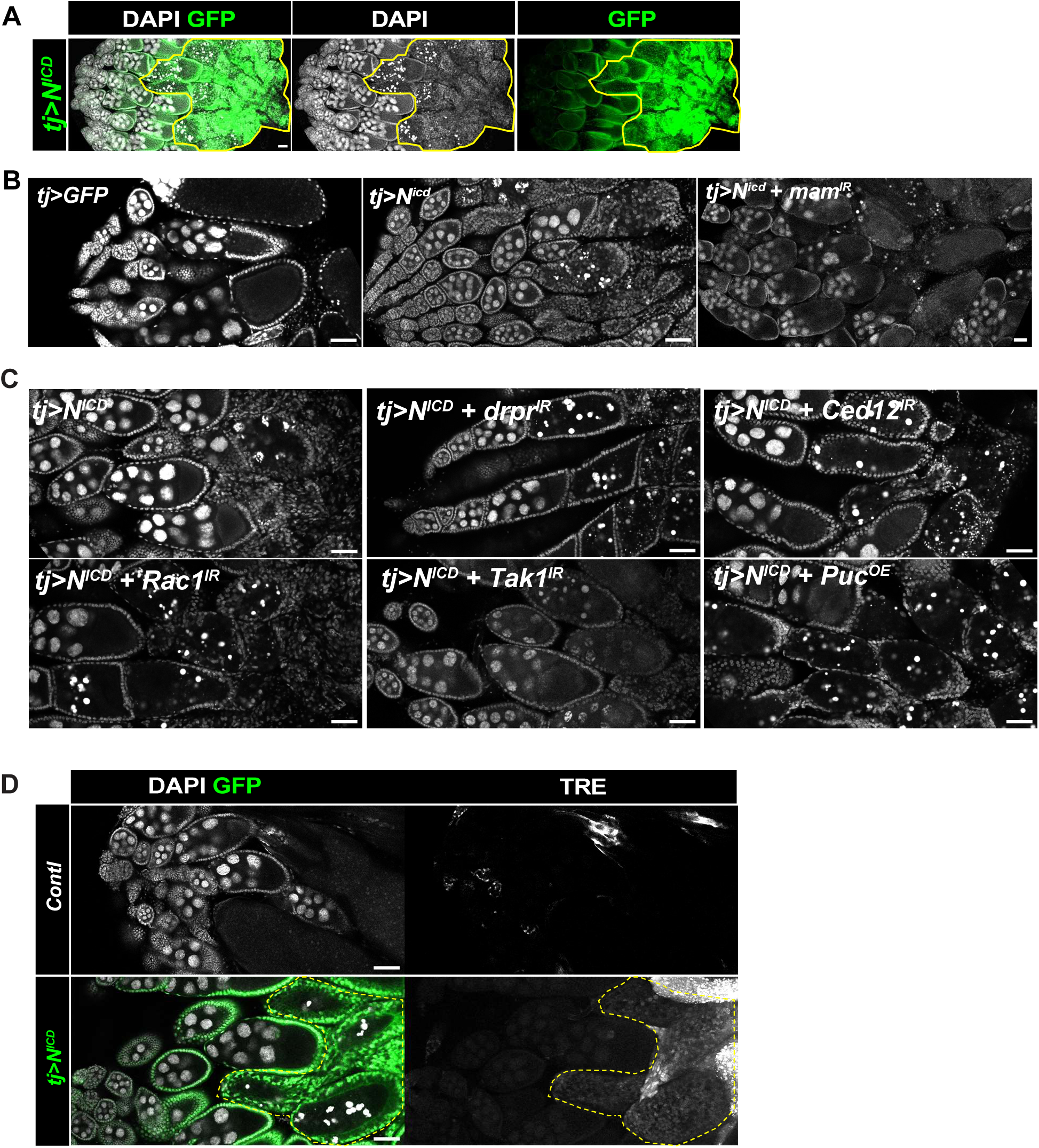
(A) Confocal images showing that overexpressing NICD in FCs resulted in large corpus luteum formation (yellow circles) and caused egg chambers to arrest development at stage 9. (B) Wide-field confocal images corresponding to Figure 3F. (C) Wide-field confocal images corresponding to Figure 4(A-C, G and H). (D) Wide-field confocal images showing JNK reporter TRE-RFP expression level in egg chambers undergoing GDAC induced by NICD-OE compared with wild-type (wt) egg chambers. DAPI was used to mark cell nuclei. Posterior is oriented to the right. Scale bars: 50 μm

**Supplementary Figure 3:**
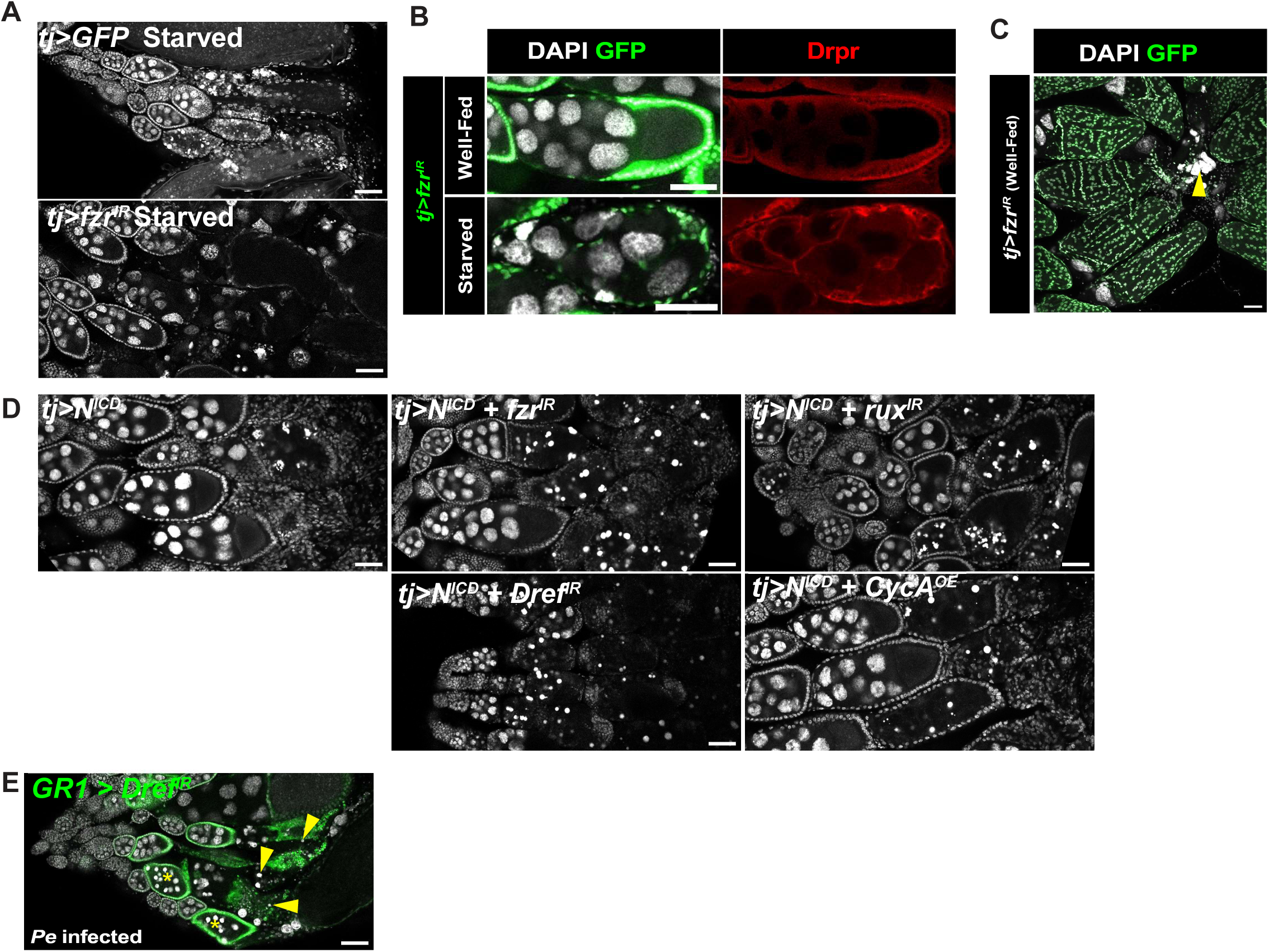
(A) Wide-field confocal images corresponding to Fig 5A-B. (B) Reducing ploidy (*fzr* knockdown) under stress conditions led to impaired GDAC. (C) Ploidy reduction in follicle cells (FCs) induced by *fzr^IR^* inhibited the developmental engulfment of nurse cell nuclei in late oogenesis, resulting in the accumulation of uncleaned germline debris in corpus luteum (yellow arrowhead). (D) Wide-field confocal images corresponding to Figure 5(D,E, G-I). (E) Knockdown of *Dref* in follicle cells resulted in inefficient GDAC under *Pe* infection (yellow arrowheads), and uncleaned nurse cell nuclear debris (DAPI staining, white). DAPI (white) was used to mark cell nuclei. Posterior is oriented to the right. Scale bars: 50 μm.

**Supplementary Figure 4:**
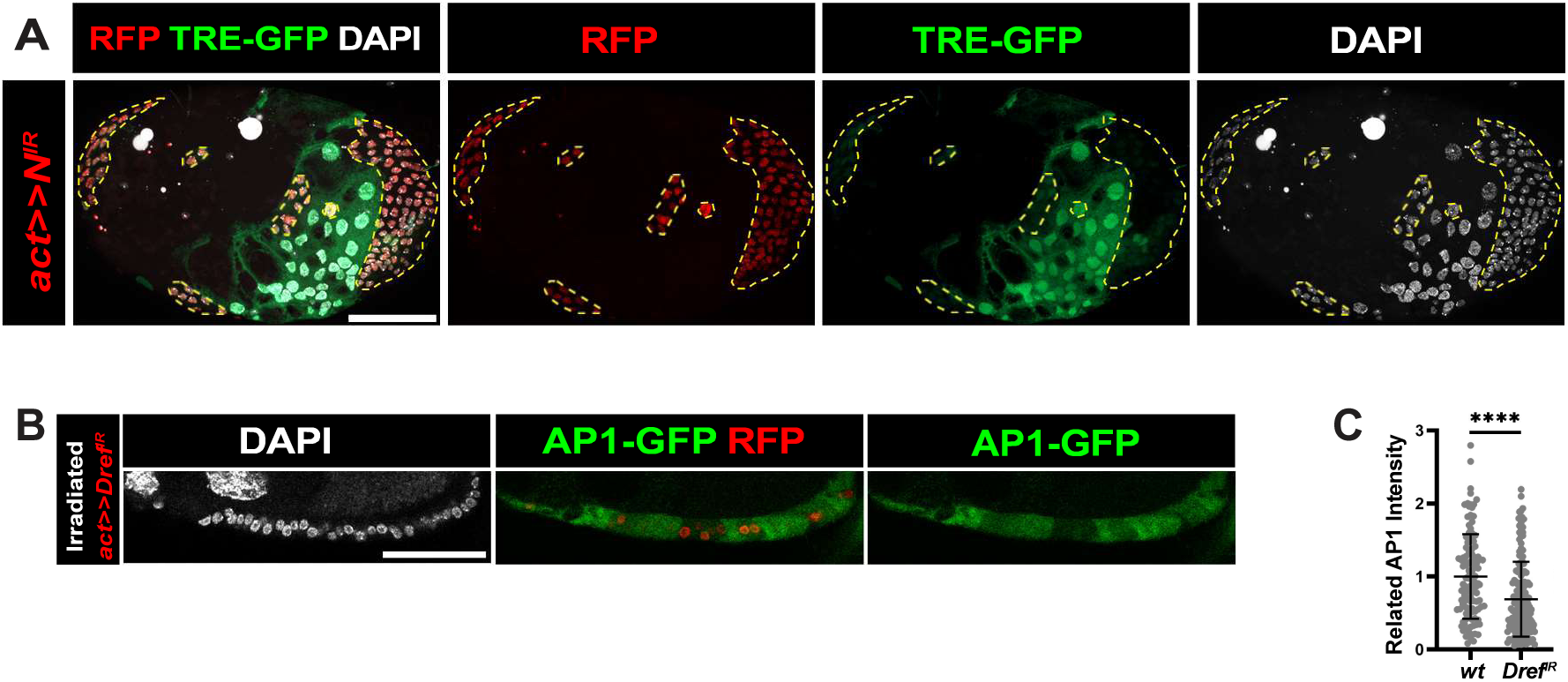
(A) Wide-field confocal images corresponding to flip-out NIR eggchamber under starvation to Fig 6A. (B) *Dref^IR^* FCs, marked by RFP, show low levels of JNK activity. (C) Graph shows (B) *Dref^IR^* FCs expressing lower AP-1 GFP. **** represents p-value ≤ 0.0001.y

---

**Supplemental Table 1:**
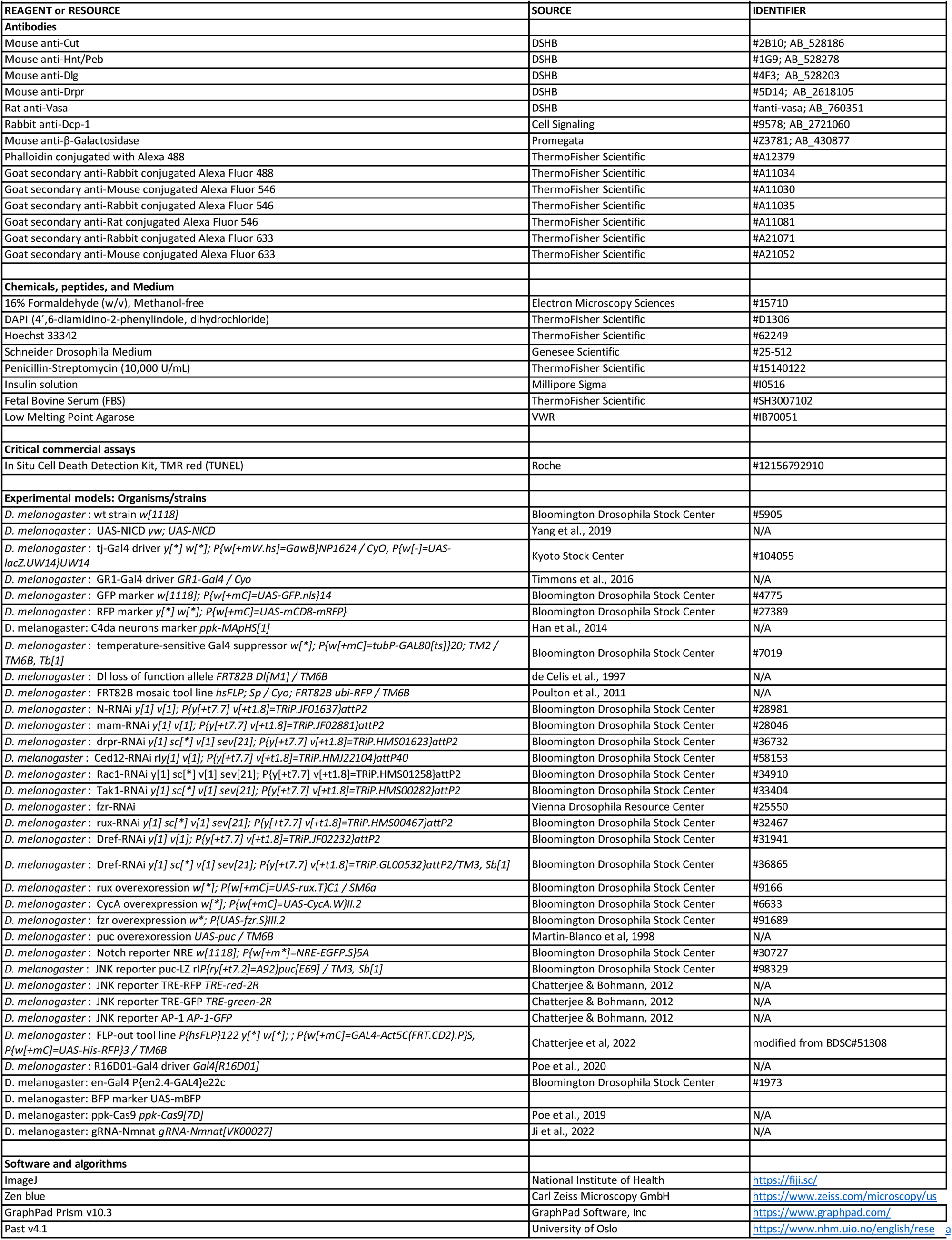
STAR Methods.

